# PCH1 regulates light, temperature, and circadian signaling as a structural component of phytochrome B-photobodies in *Arabidopsis*

**DOI:** 10.1101/566687

**Authors:** He Huang, Katrice E. McLoughlin, Maria L. Sorkin, E. Sethe Burgie, Rebecca K. Bindbeutel, Richard D. Vierstra, Dmitri A. Nusinow

## Abstract

The phytochrome (phy) family of bilin-containing photoreceptors are major regulators of plant photomorphogenesis through their unique ability to photointerconvert between a biologically inactive red light-absorbing Pr state and an active far-red light-absorbing Pfr state. While the initial steps in Pfr signaling are unclear, an early event for the phyB isoform after photoconversion is its redistribution from the cytoplasm into subnuclear foci named photobodies (PBs) that dissipate after Pfr reverts back to Pr by far-red irradiation or by temperature-dependent non-photochemical reversion. Here we present evidence that PHOTOPERIODIC CONTROL OF HYPOCOTYL 1 (PCH1) functions both as an essential structural component of phyB-containing PBs and as a direct regulator of thermal reversion that is sufficient to stabilize phyB as Pfr *in vitro.* By examining the genetic interaction between a constitutively active phyB^Y276H^-YFP allele *(YHB-YFP)* and PCH1, we show that the loss of PCH1 prevents YHB from coalescing into PBs without affecting its nuclear localization, whereas overexpression of PCH1 dramatically increases PB levels. Loss of PCH1, presumably by impacting phyB-PB assembly, compromises a number of events elicited in *YHB-YFP* plants, including their constitutive photomorphogenic phenotype, red light-regulated thermomorphogenesis, and input of phyB into the circadian clock. Conversely, elevated levels of both phyB and PCH1 generate stable, yet far red-light reversible PBs that persisted for days. Collectively, our data demonstrate that the assembly of PCH1-containing PBs is critical for phyB signaling to multiple outputs, and suggest that altering PB dynamics could be exploited to modulate plant responses to light and temperature.

**Significance:** In *Arabidopsis*, phytochrome B (phyB) perceives light and temperature signals to regulate various fundamental morphogenic processes in plants through its interconversion between its active Pfr and inactive Pr states. Upon photoconversion from Pr to Pfr, phyB forms subnuclear foci called photobodies, whose composition and molecular function(s) are unclear. We show here that the phyB-interacting protein PCH1 is a structural component of phyB-photobodies and protects Pfr from thermal reversion back to Pr thus helping maintain phyB signaling. Loss of PCH1 compromises photobody formation, which disrupts a number of downstream events including photo- and thermal perception and signaling into the circadian clock. These results demonstrate that forming PCH1-dependent phyB-photobodies is an essential step connecting light and temperature to controls on plant morphogenesis.

## Introduction

To adapt to ever-changing environments, plants integrate various external and endogenous signals throughout development and over the growing season to regulate gene expression (1). Light is an important environmental cue that triggers transcriptional reprograming and major developmental transitions. Plants sense subtle changes in light quantity, direction, duration, and color via a variety of photoreceptors that perceive and transduce signals across the visible spectrum (2-4). One major class of photoreceptors are the red/far-red light absorbing phytochromes (phys) that participate in nearly all aspects of plant growth and development, including seed germination, de-etiolation, circadian rhythms, photomorphogenesis, and foliar shade responses (5). The model plant *Arabidopsis thaliana* utilizes five phys named phyA-phyE that have both unique and overlapping functions (6), with phyB acting as the major red-light photoreceptor (3).

The phy family members possess a bilin chromophore to generate two interconvertible states, a biologically-inactive Pr state that absorbs red light (∼660 nm) and a biologically-active Pfr state that absorbs far-red light (∼730 nm) (5). In addition to the rapid red/far-red light induced photoconversion, Pfr spontaneously reverts back to Pr in a process known as thermal or dark reversion (7). Therefore, both photoconversion and thermal reversion determine the steady-state ratio Pr/Pfr, and control the extent of phytochrome signaling.

Recently, specific mutations in phyB have been identified that uncouple photoconversion from signaling. A tyrosine to histidine substitution (Tyr 276 to His) in phyB (phyB^Y276H^, hereafter referred to as *YHB)* is unresponsive to light and appears to continuously signal to downstream components, resulting in a constitutive photomorphogenic phenotype even in dark-grown seedlings (8, 9). The *YHB* allele is also sufficient in darkness to mimic constant light input into the circadian clock (10). Thus, the YHB mutant provides a tool to investigate aspects of red-light signaling downstream of phyB without activating other photoreceptors.

Another feature of phy signaling is the ability of phyB as Pfr (and other phys) to aggregate into discrete subnuclear foci called photobodies (PBs), whose connection to phy action remains enigmatic (11, 12). The accumulation of these micron-sized, bio-condensates observed using phys tagged with fluorescent polypeptides is dynamic and responsive to light, suggesting that this accretion represents an important signaling intermediate. For example, the accumulation of phyB-PBs positively correlates with several phyB-mediated events, such as the degradation of phyB targets, photoinhibition of hypocotyl elongation, and repression of leaf hyponasty (13, 14). Prolonged light exposure generates larger phyB-PBs that can prolong phyB signaling in seedlings transferred from light to darkness (14), while lowering irradiance or lowering the R-to-FR ratio (e.g. foliar shade) leads to smaller PBs and phyB evenly dispersed within the nucleoplasm (13). YHB, mimicking the active Pfr state, will readily form PBs in the absence of light. Conversely, several intragenic phyB mutations that specifically affect PB formation, but not nuclear localization, also phenocopy *phyB* loss-of-function mutants (15).

Despite their likely importance, the mechanism by which phyB-PBs assemble and the composition of phyB-PBs also remain elusive. One key factor appears to be PHOTOPERIODIC CONTROL OF HYPOCOTYL 1 (PCH1), first identified as a phyB-interacting protein that preferentially binds to Pfr (16). Subsequent studies showed that PCH1 promotes phyB-PB accumulation, and prolongs phyB activity in the dark by possibly slowing thermal reversion *in vivo* (17). Here we further analyzed the biochemical and genetic interactions between phyB and PCH1. Our results provide *in vitro* evidence that PCH1 is sufficient to slow the thermal reversion of phyB from Pfr to Pr. In addition, we found that PCH1 is required for assembly of phyB-PB *in vivo.* Using the constitutively active phyB allele YHB tagged with YFP, we show that the loss of PBs in *YHB-YFP pch1* seedlings represses the constitutive photomorphogenic phenotype of YHB, alleviates the thermo-insensitivity caused by the presence of YHB, and abolishes the YHB mediated light input into the circadian clock of dark-grown seedlings. Collectively, our work not only indicates that PCH1 is a structural component of phyB- PBs but also that forming PBs is critical for regulating multiple phyB-controlled physiological processes.

## Results

### The phyB C-terminal PASH domain is critical for interacting with PCH1

To further define how PCH1 interacts with phyB, we assayed a collection of phyB truncations for their interactions with PCH1. As illustrated in **Fig. 1*A***, phyB can be structurally divided into an N-terminal photosensing module (PSM) and the C-terminal output module (OPM), with the native phytochromobilin (PϕB) chromophore bound within the PSM. When expressed recombinantly in the presence of the synthetic chromophore phycocyanobilin (PCB), both the PSM and OPM of phyB and the full-length photoreceptor interacted with recombinant PCH1 (His_6_-PCH1-His6-Flag_3_)by *in vitro* pulldown assays done under red light (16) indicating that both domains have PCH1- binding sites **(Fig. 1*B*).** Subsequent dissections of the OPM by both *in vitro* pulldown and yeast two-hybrid assays revealed that the Period/Arnt/SIM (PASII) domain within the OPM is sufficient for this interaction, as a truncation containing only the PASII domain bound PCH1 (**Fig. 1*B*** and **1*C***).

**Fig. 1.**
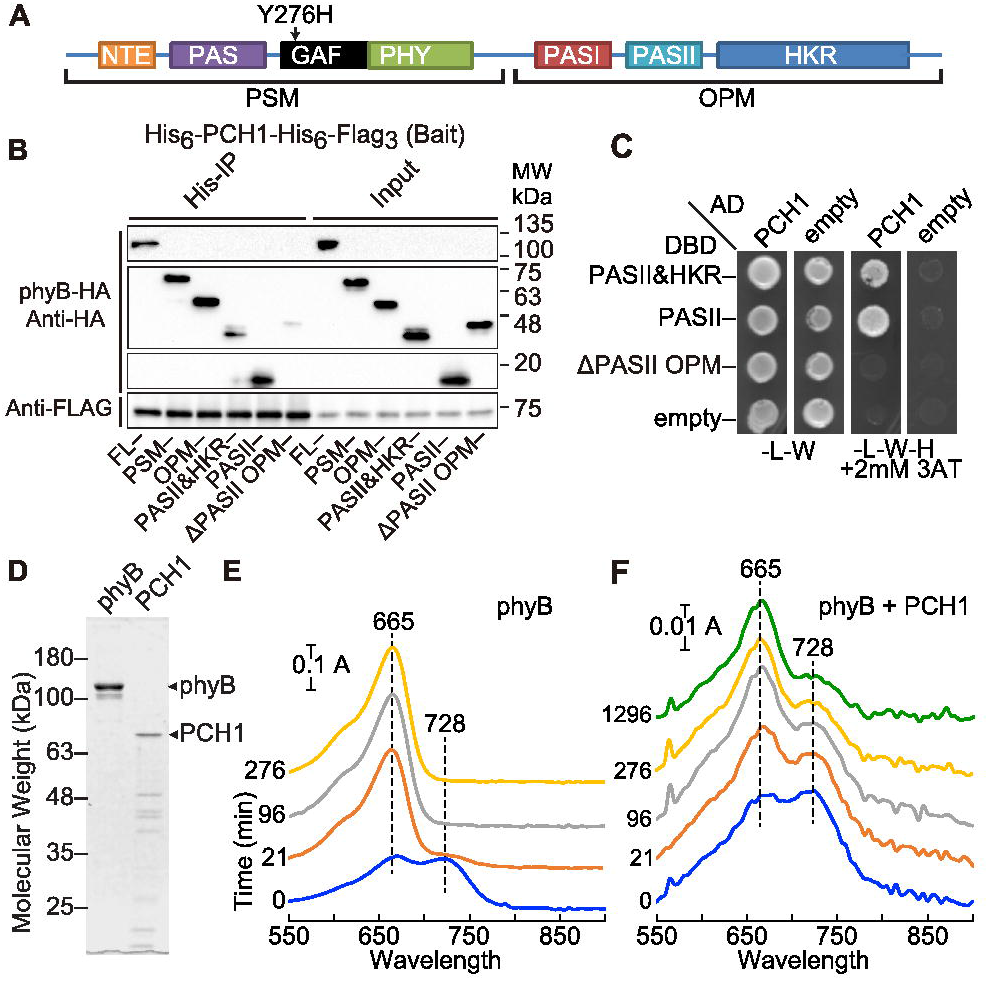
PCHl interacts with phyB C-terminal PASII domain and stabilizes phyB Pfr *in vitro*. (*A*) Illustration of full-length phyB containing N-terminal photosensing module (PSM) and C-terminal output module (OPM). NTE: the N-terminal extension; PAS: Period/Arnt/Single-Minded domain; GAF: cGMP phosphodiesterase/adenylyl cyclase/FhIA domain; PHY: Phy-specific domain; HKR: Histidine-kinase-related domain (3, 36). The tyrosine to histidine substitution (Y276H) of YHB is indicated by an arrow. (*B*) Co-immunoprecipitation (co-IP) testing interaction between phyB fragments and PCH1. Recombinant PCH1 (His_6_-PCH1-His6-Flag_3_) was used as the bait for capture (His-IP) of truncated phyB fragment preys (HA tagged). FL: full-length; APASII OPM: phyB OPM without the PASII domain. Other fragments are labeled after the domain(s) they contain. Molecular weight (MW) is labeled on the side. (*C*) Yeast two-hybrid analysis testing interaction between fragments of phyB-OPM (fused to DBD) and PCH1 (fused to AD). Vectors alone (empty) served as negative controls. (*D*) SDS-PAGE and coomassie staining showing purified recombinant PCH1 and full-length phyB. ns indicates a non-specific band of low molecular weight. (*E*) and (*F*) Absorbance spectra of Pfr to Pr thermal reversion for phyB (F) and phyB in the presence of PCH1 (F) collected at given time points under dark conditions. Differences in absorbance reading are indicated by scales. Wavelengths of absorbing peak of Pfr and Pr were labeled by dashed lines. All experiments in this figure were done at least twice with consistent results.

### PCH1 stabilizes phyB in its Pfr form *in vitro*

Enderle et al. recently showed that altering the levels of PCH1 affects the Pfr to Pr thermal reversion rate of phyB *in vivo* through an unknown mechanism (17). As the PASII interaction site within phyB that binds PCH1 overlaps with the region known to influence its thermal reversion [**Fig. 1 *B*** and ***C***, (15, 18)], we speculated that PCH1 blocks thermal reversion of phyB through direct binding. To test this hypothesis, we measured thermal reversion of purified full-length phyB assembled with the natural phytochromobilin in the presence and absence of PCH1 **(Fig. 1*D*).** Both samples photoconverted well to Pfr with red-light (blue line, **Fig. 1*E*** and ***F***). As shown in **(Fig. 1*F*)**, phyB rapidly reverted from Pfr back to Pr in the dark, with the reaction nearly complete by 21 min as judged by the loss of absorbance at 730 nm and the gain of absorbance at 660 nm for phyB. In contrast, when PCH1 was added to phyB preparations in near equimolar concentrations, thermal reversion was substantially slowed with appreciable amounts of Pfr still detected even after nearly 1,300 min in darkness **(Fig. 1*F***). In reactions containing less PCH1 relative to phyB, we could measure the impact of PCH1 on the thermal reversion rate **(SI Appendix, Fig. S1*A*).** For phyB alone, the decrease in the Pfr absorption peak fit well to two exponentials with rate constants of 0.33 and 0.038 min^−1^ **(SI Appendix, Fig. S1*B*** and **S1*C***). In the presence of PCH1, three exponentials were required to fit the data. Two rate constants 0.4 and 0.039 min^−1^ corresponded with those of free phyB, whereas the third rate constant was unique to the phyB-PCH1 sample. Its relatively small rate value (0.0013 min^−1^) implied that PCH1 reduces the thermal reversion rate of phyB by ∼30x **(SI Appendix, Fig. S1*B* and SI*C***), which agrees with *in vivo* observations [**Fig. 4** and (17)]. These results demonstrated that PCH1 was sufficient to slow the thermal reversion of phyB in a recombinant system.

### Constitutively active phyB^Y276H^ requires PCH1 to form phyB-PBs

Both *in vitro* (**Fig. 1 *E*** and ***F***) and *in vivo* spectrophotometry (17) showed that PCH1 stabilized phyB as Pfr and prolonged the Pfr-containing phyB-PBs and phyB activity. However, it was unclear whether PCH1 is a required structural component of phyB-PBs. If PCH1 function is limited to stabilizing phyB-Pfr, phyB-PBs should form in the absence of PCH1 as long as Pfr is attainable (**Fig. 2*A***, WT, a). However, if PCH1 is required for the formation of phyB- PBs, PB formation should be disrupted without PCH1, even with phyB photoconverted to Pfr (**Fig. 2*A***, WT, b). We used the constitutively active YHB mutant to investigate these possibilities (**Fig. 1*A***). If PCH1 solely regulates phyB reversion from Pfr to Pr, YHB should be epistatic to the loss of PCH1 (**Fig. 2*A***, YHB *pch1*, a). However, if PCH1 is structurally required for forming phyB-PBs, *YHB pch1* plants should have reduced PBs, despite YHB being in a constitutively active conformation **(Fig. 2*A***, *pch1* YHB, b). To test these opposing models, we crossed a *YHB-YFP phyB-9* (35S::phyB^Y276H^-YFP-Flag in the *phyB-9* null mutant) line (19) into *pch1* loss-of-function (transcript null), PCH1 overexpression *(PCH1ox)*, and *PCHlp::PCHl pchl* complementation lines (16). Since both hypotheses are based on PCH1-phyB (Pfr) interaction, we confirmed that phyB^Y276H^ binds to PCH1 by *in vitro* binding assays. While the interaction between PCH1 and wild-type phyB was light-dependent, the YHB protein bound to PCH1 in both light and darkness **(SI Appendix, Fig. S2**). This confirmed that the Y276-H mutation does not prevent PCH1 binding and that YHB assumes a conformation capable of interacting with PCH1 independent of light.

**Fig. 2.**
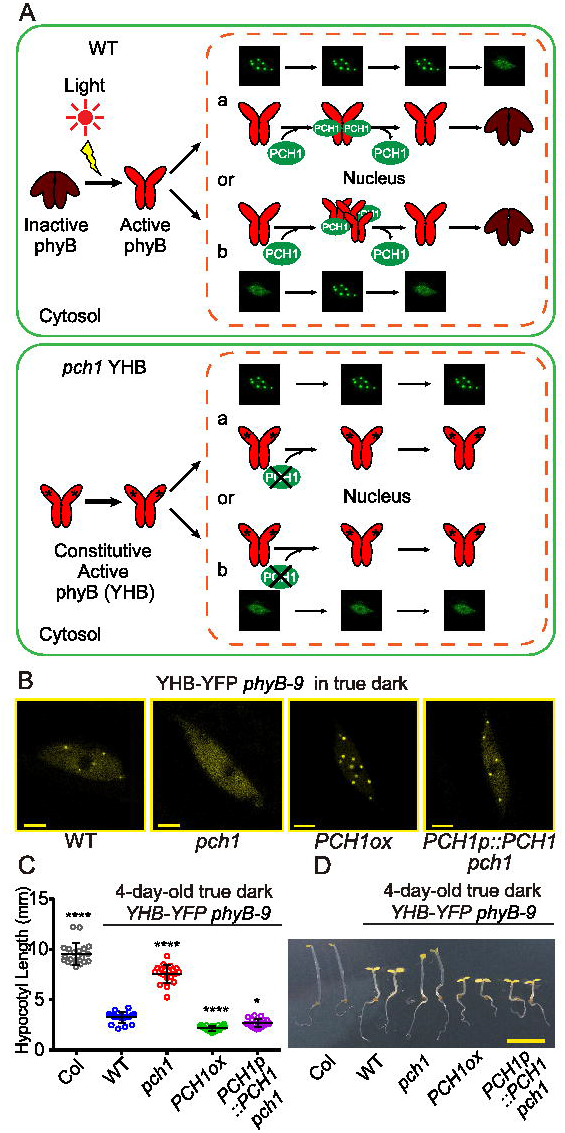
PCH1 is structurally required for the phyB-PB formation and *pchl* represses YHB constitutive photomorphogenesis. (*A*) Illustration of two possible molecular mechanisms of PCHl-mediated phyB-PB formation. phyB-GFP signals in the nucleus are used to indicate if phyB-PBs are formed (subnuclear foci vs evenly dispersed GFP signal), upper panel shows phyB-PB formation in wild type (WT) plants, lower panel presents the predicted PB phenotypes in *YHB pchl* plants according to each hypothesis. In “a”, PCH1 is a PB stabilizing factor and the constitutively active YHB will be epistatic to *pchl* and can form PBs in *pchl.* In “b”, PCH1 is a structural PB component and no PBs can be formed in the absence of PCH1. (*B*) Representative confocal images of *YHB-YFP phyB-9* in wild type (WT), *pchl, PCHlox* and *PCHlp::PCHl pchl* nuclei from elongating hypocotyl cells. Seedlings (2-day-old) grown under the true dark condition were fixed for confocal microscopy. Scale bar = 5 pm. (*C*) and (*D*) Hypocotyl elongation of 4-day-old Col-0 and *YHB-YFP phyB-9* (in various genetic background as in B) seedlings grown under the true dark condition. One-way ANOVA and Tukey’s multiple comparisons tests were conducted (n = 20). Error bars = standard deviation. * symbol indicates significantly different from WT (*, p<0.05; ****, pcO.OOOl). (*D*) shows a representative picture of seedlings in (*C*). Scale bar = 5mm. All experiments in this figure were done at least twice with consistent results.

We then grew *pch1* seedlings under a condition in which an additional far-red light pulse was applied post-germination to inactivate other phys (true dark), thus leaving YHB-YFP as the sole active phy in the dark-grown seedlings (20). In elongating hypocotyl cells, YHB-YFP maintained nuclear localization and formed subnuclear PBs, with 84% of nuclei having 2 to 5 bright foci (**Fig. 2*B*** and **SI Appendix, Fig. S3**). However, introduction of the *pch1* mutation nearly eliminated the ability of YHB-YFP to form PBs in the nucleus; here 97% of the *pch1* mutant nuclei observed harbored no PBs with the rest containing only 1 PB (**Fig. 2*B*** and **SI Appendix, Fig. S3** inset). It is noteworthy that the *pch1* mutation only affected PB formation, but not the nuclear localization of YHB-YFP, as a diffuse nuclear fluorescence signal was observed in the *pch1* backgrounds (**Fig. 2*B***). Conversely, increased PCH1 levels in *PCH1ox* seedlings yielded more YHB-PBs, with 85% of cells containing 6 or more YFP-labeled puncta per nuclei (**Fig 2*B*** and **SI Appendix, Fig. S3**). The reduced levels of PBs in *pch1* seedlings was rescued by re-introducing PCH1 into the background (**Fig. 2*B*** and **SI Appendix, Fig. S3**). Most (78%) of the *PCHlp::PCHl pchl* nuclei contained 2 to 8 YHB-YFP PBs per cell (Fig. 2*B* and **SI Appendix, Fig. S3**). Therefore, our confocal microscopy data and genetic interaction studies between *pchl* and YHB- YFP indicated that PCH1 is structurally required for phyB-PB assembly (see **Fig. 2*A***).

### *pchl* represses the constitutive photomorphogenic phenotype of YHB

Because the *pchl* mutation impairs phyB-PB formation without affecting YHB-YFP nuclear localization (**Fig. 2*B***), we could test whether phyB-PB assembly was required for phyB-mediated physiological processes. As described previously, dark-grown *YHB-YFP* seedlings exhibit constitutive photomorphogenesis as exemplified by shorter hypocotyls and open cotyledons [(19), **Fig. 2 *C*** and ***D***]. *pch1* suppressed this phenotype, with *YHB- YFP pchl* seedlings having much longer hypocotyls than those expressing YHB-YFP alone, to now approach that seen for wild-type hypocotyls in darkness (**Fig. 2*C, D*** and **SI Appendix, Fig. S4**). However, the lack of complete suppression of the effect of YHB on hypocotyl elongation by loss of PCH1 protein shows that nuclear YHB (**Fig. 2*B***) outside of photobodies still signal, albeit at severely reduced capacity. In addition, *YHB-YFP pch1* retained partially opened cotyledons, suggesting there may be separate functions for photobodies in hypocotyl elongation versus cotyledon separation, or that PCH1 could have tissue-specific roles. *PCHlp::PCHl* rescued the *pchl* mutation in the *YHB-YFP* background, while *PCH1* overexpression *(PCH1ox)* enhanced photomorphogenesis and further inhibited hypocotyl elongation in dark-grown *YHB-YFP* seedlings (**Fig. 2*C***). In contrast to our observations with YHB-YFP plants, altering PCH1 levels in wild-type, *phyB-9* or *35S::phyB-GFP phyB-9* backgrounds did not result in a constitutive photomorphogenic phenotype **(SI Appendix, Fig. S4**). Therefore, genetic interactions between *pch1* and YHB provides *in vivo* evidence that, in addition to maintaining active phyB, PCH1-mediated PB formation is essential for proper phyB signaling.

### *pchl* represses the thermomorphogenesis phenotype of YHB

Recently, phyB was discovered to provide temperature sensation and to integrate temperature with light during plant growth (21, 22). Furthermore, phyB mutants with either slowed thermal reversion or the constitutively active YHB were shown to suppress hypocotyl elongation at high temperatures (21, 22), suggesting that as a regulator of phyB, PCH1 also influences thermoresponsiveness. To assay this possibility, we measured hypocotyl elongation of seedlings grown under a short-day, high-temperature regime (SD-28°C). As expected, wild-type seedlings grown at high temperatures were significantly longer than those grown under low temperatures (SD-22°C, 28°C/22°C = 2.27 fold) (**Fig. 3*A***). By contrast, *YHB-YFP* plants were less responsive (28°C/22°C = 1.49 fold) [**Fig. 3*A*** and (21)]. The thermoresponsiveness of *YHB-YFP* plants was affected by altering PCH1 levels, as loss of PCH1 repressed the *YHB-YFP* hyposensitive phenotype (28°C/22°C = 2.22 fold) (**Fig. 3*A***). However, in *PCHlp::PCHl pchl* and *PCHlox* backgrounds, *YHB-YFP* hyposensitivity to temperature was enhanced (28°C/22°C = 1.27 and 1.36 fold, respectively) (**Fig. 3*A***). Therefore, similar to photomorphogenesis, our data imply that PCH1-dependent PBs are required for thermomorphogenesis directed by phyB.

**Fig. 3.**
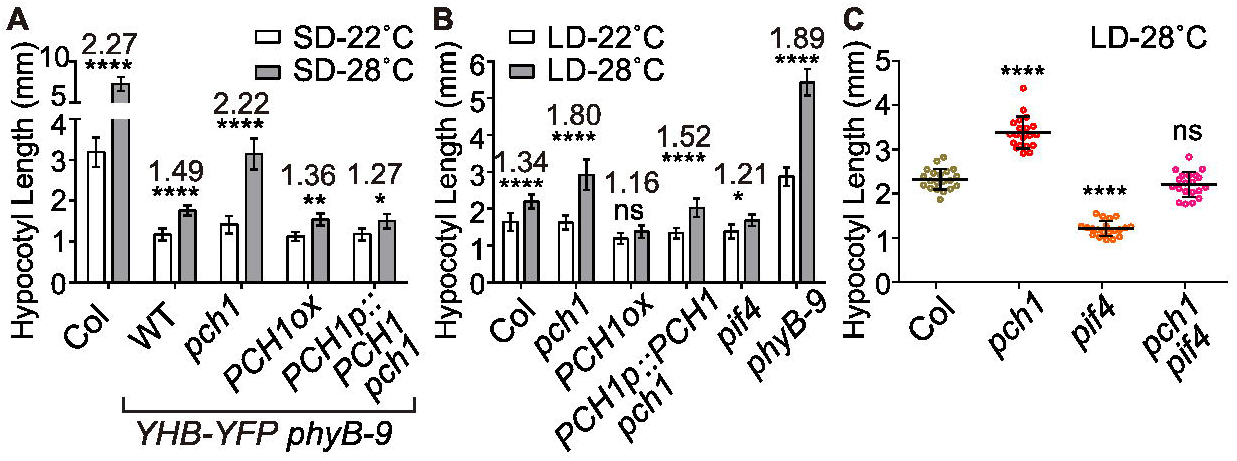
*pchl* represses YHB thermoresponses and regulates plant growth in response to elevated temperature. (*A*) Hypocotyl lengths of 4-day-old seedlings of Col-0 and *YHB-YFP phyB-9* (in WT, *pchl, PCHlox* or *PCHlp::PCHl pchl)*, under the short day photoperiod (SD) with either constant 22°C or 28°C. (S) Hypocotyl lengths of Col-0, *pchl, PCHlox, PCHlp::PCHl pchl, pif4* and *phyB-9* 4-day-old seedlings grown under the long day photoperiod (LD) with either constant 22°C or 28°C. Numbers indicate fold changes of 28 °C treated seedlings over 22°C. Two-way ANOVA and Tukey’s multiple comparisons tests were conducted comparing 28°C treatment to 22°C. (*C*) Hypocotyl lengths of Col-0, *pchl, pif4* and *pif4 pchl* 7-day-old seedlings grown under the LD-28°C condition. One-way ANOVA and Tukey’s multiple comparisons tests were conducted compared to Col-0. For all figures, n = 20, error bars = standard deviation, ns, not significant; *, p<0.05; **, p<0.01; ****, pcO.OOOl. All experiments in this figure were done at least twice with consistent results.

We also measured the response of hypocotyls from *YHB-YFP, YHB-YFP pch1, YHB-YFP PCH1ox*, and *YHB-YFP PCHlp::PCHl pchl* seedlings grown at elevated temperatures in darkness **(SI Appendix, Fig. S5**). In constant dark, all seedlings showed a similar ratio of hypocotyl length in response to elevated temperature (between 1.45-1.78 fold, 28°C/22°C). The differences observed in hypocotyl elongation between constant dark and short-day conditions may reflect the influence of other light-activated photoreceptors on hypocotyl elongation in response to temperature and light conditions.

### PCH1 directly regulates the thermoresponsive growth of plants under long days

Next, we tested whether modulating PCH1 levels in the wild-type background would be sufficient to affect the thermoresponsive growth in the absence of YHB. Arabidopsis hypocotyl elongation is inhibited in long days (LD). As shown previously (23) and in **Fig. 3*B***, elevating temperature from 22°C to 28°C suppressed the effect of the LD photoperiod and significantly promoted hypocotyl elongation of wild-type plants (28°C/22°C = 1.34 fold). Hypocotyl length of *pch1* seedlings was not significantly different from that of wild-type plants under the LD-22°C condition [**Fig. 3*B*** and (16)]. However, hypocotyl elongation of *pch1* seedlings under LDs was significantly more sensitive to high temperature (28°C/22°C = 1.80 fold) than that of wild type under the LD condition (**Fig. 3*B***). Consistent with PCH1 regulating phyB, the thermoresponsive phenotype of *pch1* seedlings resembled that of the phyB null mutant, which was also hypersensitive to high temperature (28°C/22°C = 1.89 fold) (**Fig. 3*B***). The response of *pch1* seedlings was rescued by the *PCHlp::PCHl* transgene (**Fig. 3*B***), showing that PCH1 is required for preventing extensive growth when temperatures are elevated thus counteracting the growth-inhibition seen under LDs. In contrast to *pchl* seedlings, *PCHlox* seedlings were insensitive to the temperature upshift and constitutively suppressed hypocotyl elongation under LDs regardless of the temperature (**Fig. 3*B***).

phyB-mediated light and temperature signaling converge on a group of transcription factors named PHYTOCHROME-INTERACTING FACTORS (PIFs), with PIF4 being especially influential (24). In agreement with previous reports, the *pif4* loss-of-function mutant is insensitive to high-temperature induced hypocotyl elongation when grown under constant light [(24) and **SI Appendix, Fig. S6**]. However, when grown under a LD photoperiod, *pif4* hypocotyls grew significantly longer under 28°C than under 22°C (28°C/22°C = 1.21 fold) (**Fig. 3*B***). We observed an additive genetic interaction between *pch1* and *pif4* under the LD-28°C condition, in which the *pch1* and *pif4* mutations mutually suppressed the hypocotyl phenotype caused by the other (**Fig. 3*C***). This result implied that the PCH1-mediated regulation of thermoresponsiveness was not solely dependent on PIF4 and is consistent with *PCH1ox* being insensitive and *pif4* being hyposensitive to high temperatures in long days (**Fig. 3*B***). Interestingly, when grown under continuous light, PIF4 functions downstream of PCH1 genetically, with both *pif4* and *pif4 pch1* seedlings being insensitive to high temperature **(SI Appendix, Fig. S6**). As PCH1 fine-tunes phyB signaling to prevent excessive plant growth under diel conditions (16), continuous light could mask PCH1 action and minimize the role of PCH1 in repressing the enhanced hypocotyl elongation seen at high temperatures.

### *pchl* abolishes YHB-mediated light input to the circadian clock

In Arabidopsis, the daily oscillations of light and temperature synchronize the endogenous circadian clock with the environment. This leads to the rhythmic expression of core clock genes, such as *CIRCADIAN CLOCK ASSOCIATED 1 (CCA1)*, and many clock-regulated output genes with a period of approximately 24 hrs. Such circadian rhythms can be tracked *in planta* by expressing the luciferase (LUC) reporter under the control of the *CCA1* promoter (25). Based on this reporter, plants can maintain rhythms for several days after being transferred to constant light, reflecting the free-running nature of the circadian clock. However, the cycled oscillation of LUC is considerably dampened following a switch to constant darkness (10), reflecting the need for continuous light input to maintain high amplitude oscillations of the reporter in free-running conditions.

The YHB allele mimics continuous light input and maintains high amplitude circadian rhythms in dark-grown seedlings entrained by light/dark cycles (10). Since loss of *PCH1* impairs both YHB-PB assembly and phyB-regulated light signaling, we reasoned that the loss of PBs in *YHB-YFP pch1* seedlings would also negate the input of *YHB-YFP* into the circadian oscillator. As expected, wild-type seedlings **(SI Appendix, Fig. S7*A*)** entrained to a 12 hr light:12 hr dark cycle failed to retain a high amplitude rhythm for the *CCA1::LUC* reporter after shift to constant darkness, as quantified by fast Fourier transform-nonlinear least squares and relative amplitude error analysis (26). In contrast, diurnally-entrained *YHB-YFP* seedlings maintained rhythmic *CCA1::LUC* expression following the transition to darkness for up to 144 hr (top left of **Fig. 4*A*** and **SI Appendix, Fig. S7*B*).** The *pch1* mutation eliminated this continuous rhythm, presumably by abolishing YHB-mediated light input into the clock (top right of **Fig. 4*A*** and **SI Appendix, Fig. S7*B*)**, while the *PCHlp::PCHl* transgene restored the rhythms of *YHB-YFP pch1* seedlings (middle right of **Fig. 4*A*** and **SI Appendix, Fig. S7*B*).** Overexpressing PCH1 in YHB plants increased the amplitude of the CCA1::LUC reporter as compared to wild type (bottom left of **Fig. 4*A*** and **SI Appendix, Fig. S7*B*).** Taken together, our circadian analyses indicated that although YHB is constitutively active, the PCH1-mediated formation of PBs is necessary for its influence on the amplitude of the circadian clock.

**Fig. 4.**
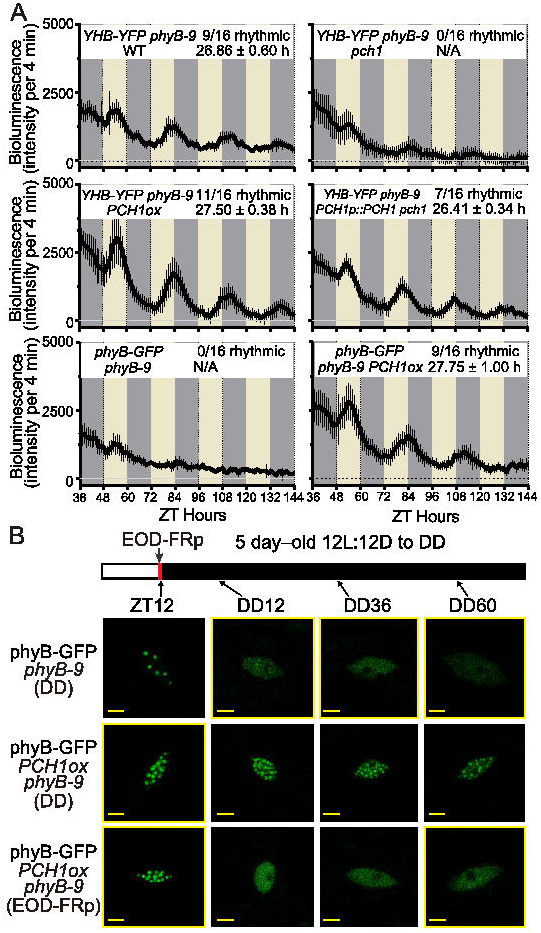
PCHl-mediated phyB-PB formation is critical for transducing light input to the circadian clock. (*A*) YHB-YFP and phyB-GFP plants (in *phyB-9)* carrying the *CCA1::LUC* reporter in different genetic backgrounds were imaged for bioluminescence under DD after 12L:12D entrainment (22°C). Each plot shows average bioluminescence of all seedlings along with standard deviation (error bars). Number of rhythmic plants (relative amplitude error (RAE) < 0.5) over all tested plants (n = 16) of each genotype are also shown. The average period length ± standard deviation of each genotype is calculated from rhythmic plants **(SI Appendix, Fig. S7*B***). Grey and darker grey blocks represent subjective light and subjective dark periods, respectively. (*B*) Representative confocal images detecting phyB-PBs in *phyB-GFP phyB-9* and *phyB-GFP PCHlox phyB-9* seedlings. 5-day-old seedlings were transferred to DD after 12L:12D entrainment and fixed at designated ZT or hours in DD for microscopy. White and black blocks above represent light and dark periods, respectively. End-of-day far-red pulse (EOD-FRp) is indicated by a red line. Scale bar = 5 pm. All experiments in this figure were done at least twice with consistent results.

### Elevated levels of both phyB and PCH1 induce persistent, reversible signaling foci

Unlike *YHB-YFP* seedlings, *phyB-9* over expressing *phyB-GFP* remain arrhythmic in the dark (bottom left of **Fig. 4*A***), consistent with wild-type phyB undergoing normal thermal reversion to Pr and thus preventing its input into the clock **(SI Appendix, Fig. S7*A*).** Similarly, overexpressing PCH1 alone does not enable plants to maintain rhythmicity after transfer to darkness *(PCH1ox*, **SI Appendix, Fig. S7*A*).** However, overexpressing both phyB and PCH1 *(phyB-GFP phyB-9 PCH1ox)* resulted in a high amplitude rhythm of the CCA1::LUC reporter (bottom right of **Fig. 4*A***).

This result suggested that the simultaneous overexpression of phyB and PCH1 could maintain a sufficient pool of phyB-PBs for extended periods when plants are transferred from a photoperiodic environment to constant darkness. To examine this notion, we assayed phyB-PB formation in *phyB-GFP phyB-9* and *phyB-GFP PCH1ox phyB-9* seedlings after periods of extended dark. After 12 hr of light treatment [Zeitgeber 12 (ZT12)], *phyB-GFP* seedlings accumulated several large PBs per nucleus (**Fig. 4*B***, top panel). Transfer to dark induced the loss of most phyB-PBs in phyB-GFP plants after 12 hr and a complete loss of large phyB-PBs by 36 hr (**Fig. 4*B***, top panel, Dark 12 and 36). By contrast, *phyB-GFP PCH1ox phyB-9* plants started with more phyB-PBs per nucleus than *phyB-GFP phyB-9* seedlings at ZT12 and retained their PBs upon extended darkness (**Fig. 4*B***, middle panel). Consistent with the ability of PCH1 to inhibit the thermal reversion of phyB **(Fig. 1*F*)** and promote PB formation (**Fig. 2*B*** and **SI Appendix, Fig. S3**), nuclear PBs were detected even after 60 hr in darkness, which suggests that maintenance of a phyB- PB in the dark is sufficient to transduce light input into the circadian clock. The accumulation of phyB-PBs in *phyB-GFP PCH1ox phyB-9* plants was reversed by a far-red pulse at the end of day (ZT12, EOD-FRp, **Fig. 4*B***, bottom panel), demonstrating that phyB-PBs in *phyB-GFP PCH1ox phyB-9* plants were still dynamic and responsive to photoconversion of phyB from Pfr back to Pr.

## Discussion

Although it has been well recognized that active phyB can form subnuclear photobodies, the structural proteins contributing to such bio-condensation remain elusive. PCH1 was recently identified as a phyB-interacting protein and possible regulator of phyB signaling. To define the function of PCH1, we used *in vitro* photochemical assays to show that PCH1 is sufficient to suppress the rate of phyB thermal reversion and extend the longevity of the Pfr signaling state, complementing a recent *in vivo* analysis (17). Furthermore, genetic interactions between *PCH1* and a constitutively active phyB allele *{YHB)* revealed that PCH1 is also an essential structural component of phyB-PB. Using *YHB-YFP pch1* plants, we in turn demonstrated the biological relevance of phyB-PBs in regulating several aspects of phyB-mediated responses, such as photo- and thermomorphogenesis. We also provided the first evidence that the assembly of phyB- PBs is critical for transducing light signals into the circadian clock.

To date, two proteins have been reported to directly regulate phyB-PB assembly: one is nuclear-localized, circadian-regulated PCH1 as discussed here and previously (16, 17), while the other is the chloroplast/nuclear dual-localized protein HEMERA (HMR) (27). By testing the genetic interaction between *YHB* and *pch1, we* discovered that PCH1 is essential for forming phyB-PBs (**Fig. 2*B*** and **SI Appendix, Fig. S3**). Similar to PCH1, a loss of function mutant of HMR also suppresses the accumulation of large phyB-PBs in light-grown phyB-GFP plants (27). However, *YHB hmr* seedlings grown in the dark plants still accumulate small PBs (27, 28), whereas PB are rarely detected in the *YHB-YFP pch1* mutant nuclei. These differences might reflect either (i) the distinct function(s) of the two proteins, (ii) our use of the true dark method to confine YHB as the lone active photoreceptor, and/or (iii) the use of fluorescent protein fusions to monitor the localization of YHB. Whether HMR and PCH1 co-regulate phyB-PB formation and whether different types of phyB-PBs exist that incorporate distinct structural components is of future interest.

Recently, the assembly of subcellular, biomolecular condensates consisting of multivalent molecules has been proposed as an important platform for effective signaling in complex cellular environments (29). Similarly, phyB-PBs could be considered as plant-specific biomolecular condensates that coalesce inside nuclei. Their formation is dynamic and responsive to light spectral quality and quantity (13, 14, 30). Not only can phyB-PBs be detected by overexpressing a fluorescent-protein-tagged phyB, they can also be seen with native photoreceptors *in planta* and be reconstituted in mammalian cells (11, 12, 31, 32). More importantly, phyB-PBs are tightly correlated with phyB functions and plant physiology (13, 14) (**Fig. 2** to **4**). As the accretion of phyB can be precisely and rapidly controlled *in vivo* by light, studying the molecular mechanisms of phyB-PB organization, dynamics, and signaling should shed light on the assembly of other biocondensates. Our results also suggest that modifying PCH1 expression could provide an avenue to fine-tune phyB signaling by directly altering thermal reversion rates and thus PB assembly and disassembly. Extending phyB activity in plants from hours to days by increasing PCH1 levels might enhance “memory” of light inputs and reduce the influence of temperature on the thermal reversion of phyB from Pfr back to Pr. We envision that manipulating PCH1 levels might help overcome challenges associated with growing crops under various photoperiod conditions, at different latitudes, and/or under the anticipated increases of temperatures caused by global warming.

## Methods

All plants used in this study are in the Col-0 ecotype *of Arabidopsis thaliana* and harbor the *CCA1::LUC* reporter (25) unless otherwise noted. *phyB-9* seeds were originally obtained from the ABRC and described previously (33). The *pch1* loss-of-function mutant (SALK_024229), *PCHlox, the PCHlp::PCHl pchl* complementation line, *phyB-GFP* transgenic plants (35S::phyB-GFP in *phyB-9), YHB-YFP* (35S::phyB^Y276H^-YFP-Flag in *phyB-9), pif4* and *pch1 pif4* were previously described (16, 19, 34). *phyB-9, phyB-GFP phyB-9, YHB-YFP phyB-9* seedlings were crossed with *pch1 CCA1::LUC, PCH1ox CCA1::LUC* and *PCHlp::PCHl pchl CCA1::LUC* seedlings. We are cognizant of a recently reported mutation in *VEN0SA4* that is present in the *phyB-9* background (35) and report *ven4* genotyping in **SI Appendix, Fig. S8**.

Detailed protocols for plant growth conditions and plasmid constructs, *in vitro* phyB thermal reversion assay, hypocotyl elongation assay and statistical analysis, fixation of seedlings and confocal microscopy, protein extraction and western blot experiments, *In vitro* co-IP/binding assay and yeast two-hybrid analysis, and luciferase-based circadian clock assays are described in **SI Appendix, SI Materials and Methods.** All primers used for cloning are listed in **SI Appendix, Table SI.**

## ACKNOWLEDGEMENTS

We thank Dr. Margaret E. Wilson for critically reading the manuscript and Jeffery Allen for technical assistance. U.S. National Science Foundation (NSF) DBI-1337680 funded acquisition of a Leica SP8-X confocal microscope. D.A.N. acknowledges support from NSF (IOS 1456796), M.L.S. was supported by NSF (DGE- 1745038). The work was supported by grants from the Molecular Biophysics Program within the NSF (MCB-1329956) and National Institute of Health (RO1 GM127892-01A1) to R.D.V.

## References

1. Scheres B & van der Putten WH (2017) The plant perceptron connects environment to development. Nature 543(7645):337–345.

2. Yu X, Liu H, Klejnot J, & Lin C (2010) The Cryptochrome Blue Light Receptors. The Arabidopsis Book/American Society of Plant Biologists 8:e0135.

3. Bae G & Choi G (2008) Decoding of light signals by plant phytochromes and their interacting proteins. Annu Rev Plant Biol 59:281–311.

4. Jenkins GI (2014) The UV-B photoreceptor UVR8: from structure to physiology. Plant Cell 26(1):21–37.

5. Li J, Li G, Wang H, & Wang Deng X (2011) Phytochrome Signaling Mechanisms. The Arabidopsis Book/American Society of Plant Biologists 9:e0148.

6. Clack T, Mathews S, & Sharrock RA (1994) The phytochrome apoprotein family in Arabidopsis is encoded by five genes: the sequences and expression of PHYD and PHYE. Plant Mol Biol 25(3):413–427.

7. Rockwell NC, Su YS, & Lagarias JC (2006) Phytochrome structure and signaling mechanisms. Annu Rev Plant Biol 57:837–858.

8. Hu W, Su YS, & Lagarias JC (2009) A light-independent allele of phytochrome B faithfully recapitulates photomorphogenic transcriptional networks. Mol Plant 2(1):166–182.

9. Su YS & Lagarias JC (2007) Light-independent phytochrome signaling mediated by dominant GAF domain tyrosine mutants of Arabidopsis phytochromes in transgenic plants. Plant Cell 19(7):2124–2139.

10. Jones MA, Hu W, Litthauer S, Lagarias JC, & Harmer SL (2015) A Constitutively Active Allele of Phytochrome B Maintains Circadian Robustness in the Absence of Light. PLANT PHYSIOLOGY 169(1):814–825.

11. Yamaguchi R, Nakamura M, Mochizuki N, Kay SA, & Nagatani A (1999) Light-dependent translocation of a phytochrome B-GFP fusion protein to the nucleus in transgenic Arabidopsis. J Cell Biol 145(3):437–445.

12. Kircher S, et al. (2002) Nucleocytoplasmic partitioning of the plant photoreceptors phytochrome A, B, C, D, and E is regulated differentially by light and exhibits a diurnal rhythm. Plant Cell 14(7):1541–1555.

13. Trupkin SA, Legris M, Buchovsky AS, Tolava Rivero MB, & Casal JJ (2014) Phytochrome B Nuclear Bodies Respond to the Low Red to Far-Red Ratio and to the Reduced Irradiance of Canopy Shade in Arabidopsis. Plant Physiol 165(4):1698–1708.

14. Van Buskirk EK, Reddy AK, Nagatani A, & Chen M (2014) Photobody Localization of Phytochrome B Is Tightly Correlated with Prolonged and Light-Dependent Inhibition of Hypocotyl Elongation in the Dark. PLANT PHYSIOLOGY 165(2):595–607.

15. Chen M, Schwab R, & Chory J (2003) Characterization of the requirements for localization of phytochrome B to nuclear bodies. Proc Natl Acad Sci USA 100(24):14493–14498.

16. Huang H, et al. (2016) PCH1 integrates circadian and light-signaling pathways to control photoperiod-responsive growth in Arabidopsis. eLife 5:e13292.

17. Enderle B, et al. (2017) PCH1 and PCHL promote photomorphogenesis in plants by controlling phytochrome B dark reversion. Nat Commun 8(1):2221.

18. Elich TD & Chory J (1997) Biochemical Characterization of Arabidopsis Wild-Type and Mutant Phytochrome B Holoproteins. The Plant Cell 9(12):2271.

19. Feng CM, Qiu Y, Van Buskirk EK, Yang EJ, & Chen M (2014) Light-regulated gene repositioning in Arabidopsis. Nat Commun 5:3027.

20. Leivar P, et al. (2008) Multiple phytochrome-interacting bHLH transcription factors repress premature seedling photomorphogenesis in darkness. Curr Biol 18(23):1815–1823.

21. Jung JH, et al. (2016) Phytochromes function as thermosensors in Arabidopsis. Science 354(6314):886–889.

22. Legris M, et al. (2016) Phytochrome B integrates light and temperature signals in Arabidopsis. Science 354(6314):897–900.

23. Nomoto Y, et al. (2012) A circadian clock- and PIF4-mediated double coincidence mechanism is implicated in the thermosensitive photoperiodic control of plant architectures in Arabidopsis thaliana. Plant Cell Physiol 53(11):1965–1973.

24. Koini MA, et al. (2009) High temperature-mediated adaptations in plant architecture require the bHLH transcription factor PIF4. Curr Biol 19(5):408–413.

25. Pruneda-Paz JL, Breton G, Para A, & Kay SA (2009) A functional genomics approach reveals CHE as a component of the Arabidopsis circadian clock. Science 323(5920):1481–1485.

26. Plautz JD, et al. (1997) Quantitative Analysis of Drosophila period Gene Transcription in Living Animals. Journal of Biological Rhythms 12(3):204–217.

27. Chen M, et al. (2010) Arabidopsis HEMERA/pTAC12 initiates photomorphogenesis by phytochromes. Cell 141(7):1230–1240.

28. Galvao RM, etal. (2012) Photoactivated phytochromes interact with HEMERA and promote its accumulation to establish photomorphogenesis in Arabidopsis. Genes & Development 26(16):1851–1863.

29. Banani SF, Lee HO, Hyman AA, & Rosen MK (2017) Biomolecular condensates: organizers of cellular biochemistry. Nat Rev Mol Cell Biol.

30. Klose C, Viczián A, Kircher S, Schäfer E, & Nagy F (2015) Molecular mechanisms for mediating light-dependent nucleo/cytoplasmic partitioning of phytochrome photoreceptors. New Phytol 206(3):965–971.

31. Hisada A, et al. (2000) Light-Induced Nuclear Translocation of Endogenous Pea Phytochrome A Visualized by Immunocytochemical Procedures. The Plant Cell 12(7):1063–1078.

32. Beyer HM, et al. (2015) Red Light-Regulated Reversible Nuclear Localization of Proteins in Mammalian Cells and Zebrafish. ACS Synth Biol 4(9):951–958.

33. Reed JW, Elumalai RP, & Chory JC (1998) Suppressors of an Arabidopsis thaliana phyB mutation identify genes that control light signaling and hypocotyl elongation. Genetics 148(3):1295–1310.

34. Nusinow DA, et al. (2011) The ELF4-ELF3-LUX complex links the circadian clock to diurnal control of hypocotyl growth. Nature 475(7356):398–402.

35. Yoshida Y, et al. (2018) The Arabidopsis phyB-9 Mutant Has a Second-Site Mutation in the VENOSA4 Gene That Alters Chloroplast Size, Photosynthetic Traits, and Leaf Growth. Plant Physiol 178(1):3–6.

36. Burgie ES & Vierstra RD (2014) Phytochromes: an atomic perspective on photoactivation and signaling. Plant Cell 26(12):4568–4583.

