## Supplemental data for "PCH1 regulates light, temperature, and circadian signaling as a structural component of phytochrome B-photobodies in *Arabidopsis*"

### Supplementary Information

**Supplementary Materials and Methods.** Detailed materials and methods used in this paper.

#### Plant Growth Conditions

Seeds were surface sterilized by 20% bleach and plated on 1/2X Murashige and Skoog medium with 0.8% agar and 1% sucrose (w/v) unless noted. After stratification for 2 to 4 days in darkness at 4°C, plates were placed in chambers for 2 to 5 days. For light-grown seedlings, plants were supplied with white light (WL, 80 $\mu mol \cdot m^{-2} \cdot s^{-1}$) under various photoperiods, including constant light, long day, 12L:12D and short day conditions (Light : Dark= 24 : 0, 16 : 8, 12 : 12 and 8 : 16 hours, respectively). For the true dark treatment, stratified seeds were treated in following steps: red light (40 $\mu mol \cdot m^{-2} \cdot s^{-1}$) for 10 min to initiate germination, then dark for 3 hours, a far-red pulse (FR, 730nm, 30 $\mu mol \cdot m^{-2} \cdot s^{-1}$) for 10 min and followed by continuous darkness for 2~4 days as indicated. For protein extraction, seedlings were grown on sterilized qualitative filter paper (Whatman, Maidstone, United Kingdom) placed on growth media for 5 days at 22 ˚C under the short day condition. The temperature for all growing conditions was set at constant 22 ˚C unless noted. For 28˚C treatment, seedlings were first grown under 22 ˚C for 24 hours to germinate before being transferred to the higher temperature.

#### Cloning of phyB constructs

cDNA of full length phyB and phyB fragments (without or with a stop codon) was cloned from the pCMX-PL2-phyB-HA construct described previously (1, 2). cDNA encoding a phyB OPM without PASII domain (phyB-OPM-ΔPASII) was made by two sequential PCRs with overlapping primers that skip the PASII domain. First, both phyB-PASI and phyB-HKR fragments were cloned using designated overlapping primers with fewer cycles of the PCR program. Then both PCR products were mixed, diluted and served as the template to clone phyB-OPM- ΔPASII using the forward primer of phyB-PASI and the reverse primer of phyB-HKR. All final PCR products were TOPO cloned into the pENTR/D-TOPO vector (ThermoFisher Scientific) and were transformed into MAX Efficiency™ Stbl2™ Competent Cells (ThermoFisher Scientific). The pENTR-phyB$^{WT}$-nonstop was then used as the template for site-mutagenesis to make pENTR-phyB$^{YH}$-

nonstop via overlapping PCR. The sequence of all pENTR constructs were verified before use.

To clone phyB$^{WT}$, phyB$^{YH}$ and phyB fragments into the pCMX vector for *in vitro* transcription/translation (IVT), we first made a gateway compatible destination vector pCMX-GW-HA. pCMX-PL2-phyB-HA was digested with KpnI (upstream of phyB) and NheI (downstream of HA) and the HA tag was re-introduced along with mutating the NheI by two partially overlapping primers pDAN0820 and pDAN0821 and In-Fusion® HD cloning (Clontech, TaKaRa)(**Table S1**). Next, we inserted an in-frame attR1-ccdB-attR2 cassette upstream of HA into the KpnI site. This cassette was cloned from the gateway-compatible destination vector pB7HFC (3) with primers pDAN0830 and pDAN0831 (**Table S1**) and then cloned into the KpnI linearized pCMX-PL2-HA vector using In-Fusion® HD cloning (Clontech, TaKaRa). The design of the forward primer abolishes the KpnI site after infusion cloning. All pENTR constructs were then gateway cloned (LR reaction, Invitrogen) into the pCMX-PL2-GW-HA vector.

**Recombinant expression and purification of full-length phyB**

Purification of recombinant full-length phyB with native phytochromobilin (PΦB) chromophore was as described (4). In brief, phyB cell pellets were thawed and sonicated with a threefold excess of lysis buffer (10% glycerol, 20 mM HEPES-NaOH (pH 7.8), 500 mM NaCl, 1 mM 2-merceptoethanol, 0.05% Tween 20, 1 mM phenylmethanesulfonyl fluoride, Pierce Protease Inhibitor Tablets (Thermo Fisher Scientific, 1 tablet per liter buffer), and 30mM imidazole) to disrupt the cells. The cell lysate was clarified by centrifugation at 35,000xg for 30 min at 4 °C. The supernatant was then loaded onto a $Ni^{2+}$-nitrilotriacetic acid (Ni-NTA) column (Qiagen) equilibrated in lysis buffer containing 30 mM imidazole. The column was washed with two column volumes of the 30 mM imidazole lysis buffer, and the bound proteins were eluted with lysis buffer containing 300 mM imidazole. Fractions containing phyB were pooled and exchanged into an anion exchange buffer (10% glycerol, 20 mM HEPES-NaOH (pH 7.8), 10 mM 2-merceptoethanol, 1mM PMSF, Pierce Protease Inhibitor Tablets (Thermo Fisher Scientific, 1 tablet per liter buffer) and 20 mM NaCl) by ultrafiltration (Amicon Ultra 15 filters), and loaded onto a HiTrap Q Sepharose HP column (GE) that was pre-equilibrated with the same buffer. Protein was eluted using a linear gradient of 20-500 mM NaCl, and fractions containing phyB were pooled and exchanged into lysis buffer using ultrafiltration (Amicon Ultra 15 filters). phyB was further enriched with a Ni-IMAC column (GE), using a 30-140 mM linear imidazole gradient in lysis buffer. Fractions containing full length phyB were pooled and incubated with 1 mg of 6His-TEV protease for three hours at 4 °C. To capture untagged phyB, protein was back-exchanged into lysis buffer containing 30 mM imidazole and passed through a Ni-IMAC column that was pre-equilibrated with the same buffer. The purified phyB sample was concentrated and passed through a Superdex 200 column (GE) in order to exchange into the appropriate buffer and remove additional contaminants. To reduce issues with PhyB degradation the protein was often flash frozen directly in liquid nitrogen between concentration steps as ~30 μl droplets.

**Monitoring phyB dark/thermal reversion using UV/Vis spectroscopy**

All spectroscopic experiments were conducted at 25 ˚C in 50 mM HEPES-KOH (pH 7.8), 1 mM $Na_4EDTA$, 10 mM 2-mercaptoethanol, and 150 mM KCl using a Cary 60 spectrophotometer (Agilent). For Pfr to Pr thermal reversion experiments, full-length (FL) phyB was converted to the Pfr form using an LED with peak absorption at 630 nm until a steady state had been reached. After removing the light source, spectra between 550 and 900 nm were collected in a time-resolved manner. For thermal reversion experiments conducted in the presence of PCH1, FL phyB and PCH1 were either combined in a 1:2 molar ratio or with phyB in molar excess. To simplify observation of the amount of Pfr state remaining, background correction was conducted by subtraction of the buffer absorption spectrum from the time resolved spectra of thermal reversion. With PCH1 in molar excess, the PCH1-phyB mixture was prone to light-scattering from aggregates. To correct for its effects on phyB chromophore absorption, the absorption due to scattering was estimated by fitting the buffer corrected baseline using absorption values at 550-600, and 830-900 nm to the equation: Baseline = $A \bullet Abs^{-4} + B$. The calculated coefficients A and B from the best fit were then used to calculate a correction for scattering that was subtracted from the observed spectra. Since scatter was variable over time, each time point was corrected individually. All calculations were conducted using R as previously described (4). Global fits of the Pfr absorption peak as a function of time were conducted using the equation $Abs = Abs_{\infty} + \sum_{i=1}^{n}(\Delta Abs_i \cdot e^{-k_i t})$, where Abs is absorbance, $Abs_{\infty}$ is the absorbance at the completion of the reaction, $\Delta Abs_i$ and $k_i$ are the amplitude and rate constant of the ith exponential and t is the time. For free PhyB n=2, and for PhyB in the presence of PCH1 n=3.

**Hypocotyl elongation assays and statistical analysis**

4-day-old or 7-day-old seedlings grown under different light/dark and temperature conditions were arrayed and photographed with a ruler for measuring hypocotyl length using the Image J software (NIH, Bethesda, Maryland). Statistical analyses (one-way or two-way ANOVA analysis with Tukey's test for multiple comparisons) for all experiments were performed using PRISM software (Graphpad, La Jolla, California, version 6.05, Graphpad.com).

**Fixation of seedlings and confocal microscopy**

For true-dark treated seedlings, the fixation solution consists of 1/2X microtubule-stabilizing (MTSB) buffer (final concentration of 25mM PIPES, 2.5mM $MgCl_2$, 2.5mM EGTA) supplemented with 4% paraformaldehyde. 2-day-old true-dark treated seedlings were fixed by soaking in fixation solution for 2 hours while still in dark growth chamber, then were rinsed twice with 1/2X MTSB buffer in dark and stored in darkness at 4°C before confocal microscopy.

For the 12L:12D to DD circadian clock assay, 12L:12D entrained seedlings of *phyB-GFP phyB-9* and *phyB-GFP PCH1ox phyB-9* were first fixed at the end of daylight (Zeitgeber 12, ZT12) on the 5th day using a filtration-based fixation method described previously (5). After transferred to continuous darkness, *phyB-GFP phyB-9* and *phyB-GFP PCH1ox phyB-9* seedlings were then sampled and fixed in dark at Dark 12 (12 hours in dark), Dark 36 and Dark 60. End-of-day far-red pulse (EOD-FRp) treatment (10min far-red light at 30 $\mu mol \cdot m^{-2} \cdot s^{-1}$) was done to 12 hr light :12hr dark entrained *phyB-GFP PCH1ox phyB-9* seedlings on 5th day, 10min before ZT12. Then seedlings were protected from any light and immediately fixed in darkness for ZT12 samples.

Confocal microscopy was performed with a Leica TCS SP8 confocal laser scanning microscope and an HC PL APO CS2 40x/1.20 WATER objective lens (Leica Microsystems, Mannheim, Germany). Light source is provided by the White Light Laser (WLL), while all emission fluorescence signals were detected by the HyD™ detector. GFP fluorescence was monitored by a 490-535 nm band emission and a 470 nm excitation line of an Ar laser. YFP fluorescence was monitored by a 525-570 nm band emission and a 514 nm excitation line of an Ar laser. Line average was set as 16 to reduce noise.

**Protein extraction, western blots analysis and *in vitro* co-IP/binding assay**

For *in vitro* co-IP/binding assay, the recombinant protein $His_6$-PCH1-$His_6$-$Flag_3$ or $His_6$-YFP-$His_6$-$Flag_3$ were used and have been described previously (1). The $phyB^{WT}$-HA, $phyB^{YH}$-HA and phyB fragments-HA preys were transcribed/translated *in vitro* (IVT) using the pCMX-$phyB^{WT}$/$phyB^{YH}$/phyB fragments-HA constructs and the TNT® T7 Quick Coupled Transcription/Translation System (Promega, Madison, Wisconsin) as instructed. Recombinant bait proteins and IVT preys were first resuspended in 500 µl PBS (150mM NaCl, 10mM Na2HPO4, 1.8mM KH2PO4, 0.1% Triton X-100) supplemented with 1 mM PMSF, 1x protease inhibitor cocktail (Roche, Pleasanton, California), 1x phosphatase Inhibitors II & III (Sigma), 50 µM MG132 (Peptides International, Louisville, Kentucky) and 20 µM phycocyanobilin (PCB). Then baits and preys were incubated for 1 hour at 15°C under either constant red (75 $\mu mol \cdot m^{-2} \cdot s^{-1}$) or dark conditions. Dynabeads™ His-Tag Isolation and Pulldown beads (Invitrogen™) were used for immunoprecipitating each sample (incubated for 20 min at 15°C), followed by washes with PBS buffer five times. Samples of input and His-IP were denatured in SDS-PAGE loading dye, separated by SDS-PAGE, transferred to nitrocellulose membrane, and detected by anti-FLAG-M2-HRP antibody (A8592, Sigma Aldrich, St Louis, Missouri) at 1:10,000 (for PCH1) and co-detected by anti-HA-HRP (Cat. 12013819001, Roche, Pleasanton, California) diluted into phosphate-buffered saline (PBS) + 0.1% Tween at 1:2000 (for interacting phyB/phyB-fragments). After washing, chemiluminescent substrate (Pierce™ ECL Western Blotting Substrate, ThermoFisher Scientific) was applied to the nitrocellulose membrane for detection.

**Yeast two-hybrid assay**

Yeast two-hybrid assays were carried out as previously described (1). In brief, the DNA binding domain (DBD) fused phyB OPM fragments or activating domain (AD) fused PCH1 constructs were transformed using the Li-Ac transformation protocol (Clontech) into *Saccharomyces cerevisiae* strain Y187 (MATα) and the AH109 (MATa), respectively. After mating two yeast strains, diploid yeast was grown on the CSM –Leu –Trp medium, and protein-protein interaction was tested in diploid yeast by replica plating on CSM –Leu

–Trp –His media supplemented with extra Adenine (30mg/L final concentration) and 2 mM 3-Amino-1,2,4-triazole (3AT). Pictures were taken after 4-day incubation at 30˚C.

**Circadian Assays in *Arabidopsis***

For circadian assays, a luciferase-based assay using the *CCA1::LUC* reporter was monitored as described previously (3). Seedlings were first grown on 1/2x MS+1% sucrose media for 6 to 8 days under the 12hr light: 12hr dark condition. Entrained seedlings were then transferred to fresh 1/2x MS without sucrose media and sprayed with sterile 5 mM luciferin (Gold Biotechnology, St. Louis, MO) prepared in 0.1% (v/v) Triton X-100 solution. Sprayed seedlings were transferred to constant darkness (22°C) and images were taken each hour over 5 to 6 days using an ultra-cooled CCD camera (Pixis 1024B, Princeton Instruments) driven by Micro-Manager software (6, 7). We used a four-minute exposure time to capture bioluminescence signal from the luciferase reporter. The images were processed by Metamorph software (Molecular Devices, Sunnyvale, CA), and rhythms determined by fast Fourier transformed non-linear least squares (FFT-NLLS) (8) after background subtraction using the interface provided by the Biological Rhythms Analysis Software System 3.0 (BRASS) available at http://www.amillar.org. FFT-NLLS analysis was conducted from ZT20- ZT144 (used ZT36 to 144 to plot) to measure period and relative amplitude error (RAE) values. Plants with a RAE value ≤ 0.5 are considered rhythmic (9).

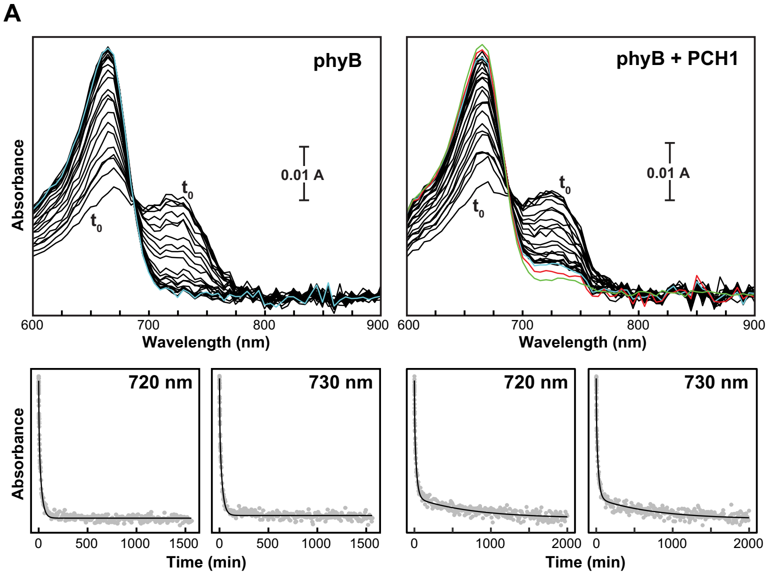


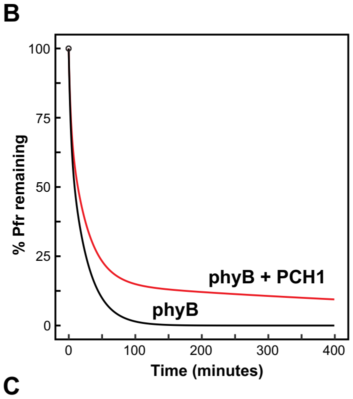


C

**Rate Constants for Pfr Thermal Reversion +/- PCH1**

| ($min^{-1}$) | phyB | phyB + PCH1[a] |
|---|---|---|
| k1 | 3.3 (±0.2) x $10^{-1}$ | 4.0 (±0.3) x $10^{-1}$ |
| k2 | 3.8 (±0.1) x $10^{-2}$ | 3.9 (±0.1) x $10^{-2}$ |
| k3 | | 1.3 (±0.1) x $10^{-3}$ |

[a] phyB was added in molar excess

**Fig. S1.** PCH1 inhibits thermal reversion of phyB from Pfr to Pr. (A) phyB in molar excess was incubated without or with PCH1 and irradiated with saturating red light until steady state was achieved (indicated by the t=0 spectra). The samples were then incubated in darkness and spectra were collected periodically as indicated by the black lines. Spectra collected at 280, 2000, and 7000 min are colored cyan, red, and green, respectively. Data from the 715-745 nm region were fit globally as a function of time to a double (phyB) or a triple exponential function (phyB + PCH1). Representative fits for 720 and 730 nm are shown below. (B,C) Simulation of the effect of PCH1 on phyB thermal reversion (B) calculated from the rate constants presented in (C) and reflect the reaction amplitudes calculated from data in (A) and (B), where phyB is in molar excess.

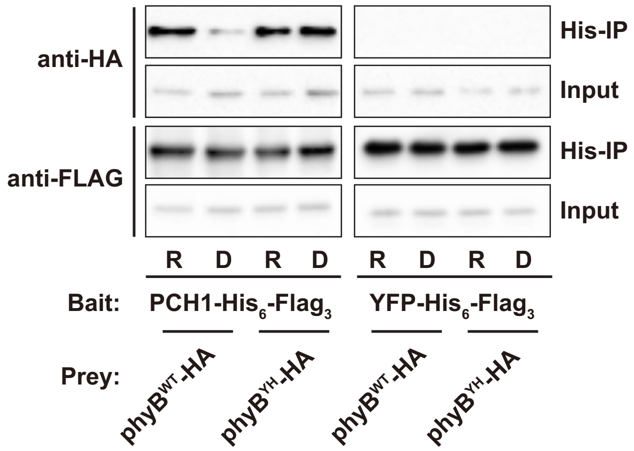


**Fig. S2.** Co-IP testing interaction between phyB$^{YH}$ and PCH1 under constant red light (R) or dark (D) conditions.

Recombinant PCH1 (His$_6$-PCH1-His$_6$-Flag$_3$) was used as the bait to capture (His-IP) *in-vitro* transcribed/translated phyB$^{WT}$-HA or phyB$^{YH}$-HA preys. Samples of input and His-IP were detected by anti-FLAG (for PCH1) and co-detected by anti-HA (for interacting phyB$^{YH}$). Recombinant YFP (His$_6$-YFP-His$_6$-Flag$_3$) protein was used as a negative control. This assay was done twice with consistent results.

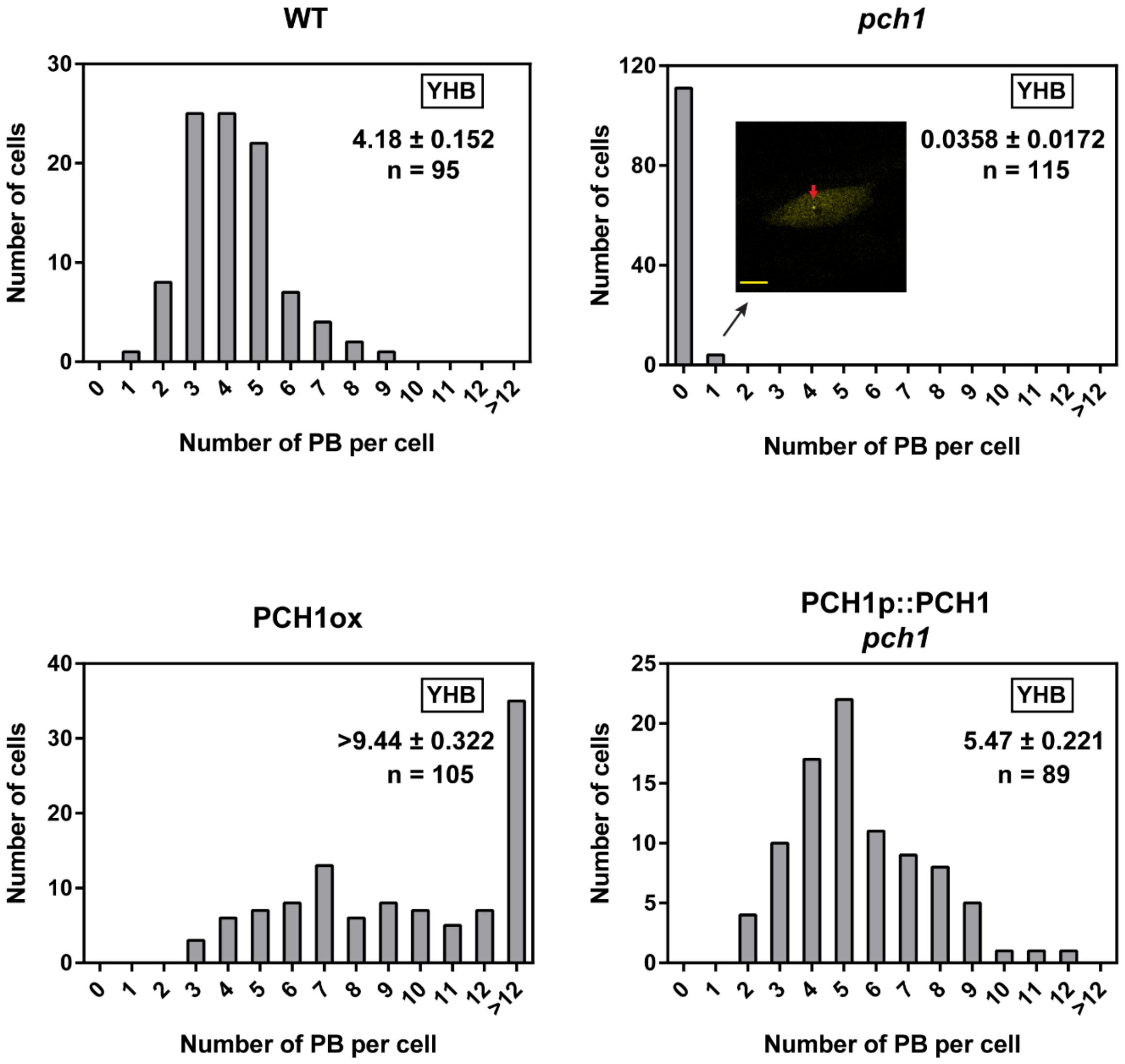


**Fig. S3.** Quantification of phyB-PBs in *YHB-YFP phyB-9* seedlings with altered PCH1 levels under the true dark condition.

Quantification was performed in *YHB-YFP phyB-9* seedlings in the WT, *pch1, PCH1ox* and *PCH1p::PCH1 pch1* backgrounds. The average number of PBs per cell for each genotype is shown with standard error of the mean. For *YHB-YFP PCH1ox phyB-9*, cells containing more than 12 PBs were binned together due to the inability to accurately count more than 13 large PBs per nucleus. Thus, the average number of PBs in *YHB-YFP PCH1ox phyB-9* is calculated by limiting the maximal number of PBs to 13. Therefore, the actual average number of PBs per cell in *YHB-YFP PCH1ox phyB-9* is likely larger than the calculated number in the figure and is represented by a "≥". An inset of the *YHB-YFP phyB-9 pch1* graph shows a representative picture of cells carrying one small PB (red arrow) with scale bar = 5 µm. Figure represents one of two independent experiments which have consistent results.

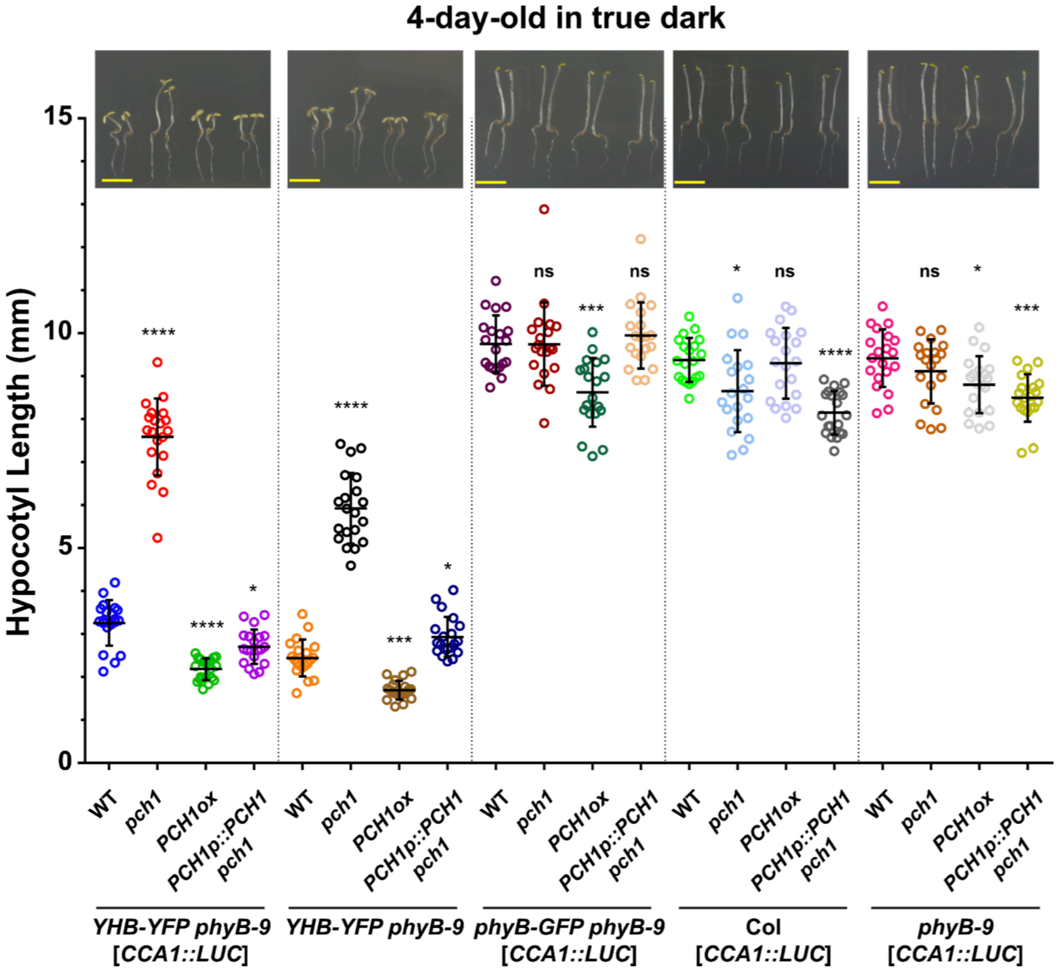


**Fig. S4.** Hypocotyl elongation of *YHB-YFP phyB-9* and *phyB-GFP phyB-9* dark-grown seedlings.

Figure shows hypocotyl lengths of true-dark-treated, 4-day-old *YHB-YFP phyB-9*, *phyB-GFP phyB-9*, Col-0 and *phyB-9* mutant backgrounds with altered PCH1 levels (WT, *pch1*, *PCH1ox* and *PCH1p::PCH1 pch1*). The *YHB-YFP phyB-9* set without the *CCA1::LUC* reporter were also included. Representative pictures of seedlings were shown above the graph with a scale bar representing 5mm. The experiment in this figure was done twice with consistent results.

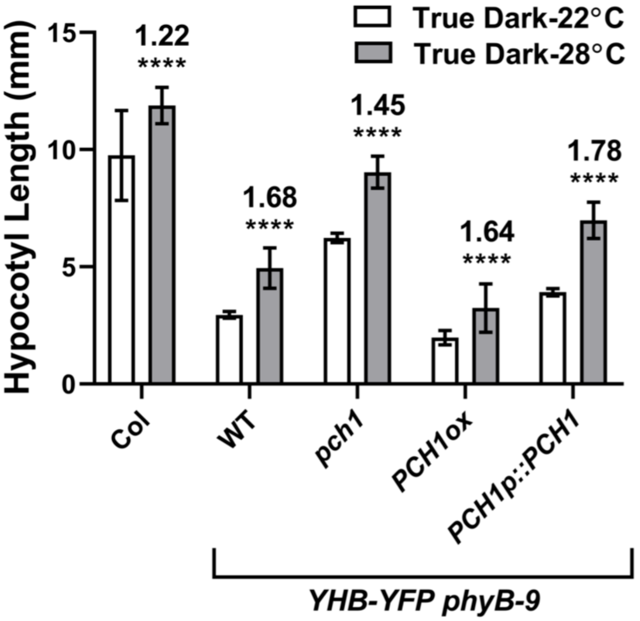


**Fig. S5.** Hypocotyl responses of *YHB-YFP phyB-9* seedlings grown in darkness at 22˚C or 28˚C.

Hypocotyl lengths of true-dark-treated, 4-day-old *YHB-YFP phyB-9* seedlings with altered PCH1 levels (WT, *pch1*, *PCH1ox* and *PCH1p::PCH1 pch1,* without the *CCA1::LUC* reporter) grown at 22˚C or 28˚C. Col-0 was included as a control. Data are averaged from 3 biological replicates where n=20 for each replicate. Two-way ANOVA and Tukey's multiple comparisons tests were conducted comparing 22˚C and 28˚C. Error bars = standard deviation. * symbol indicates hypocotyl length was significantly different between temperature treatments (*, $p<0.05$; ****, $p<0.0001$). Values above the * symbol indicate fold changes of 28˚C hypocotyl lengths over 22˚C.

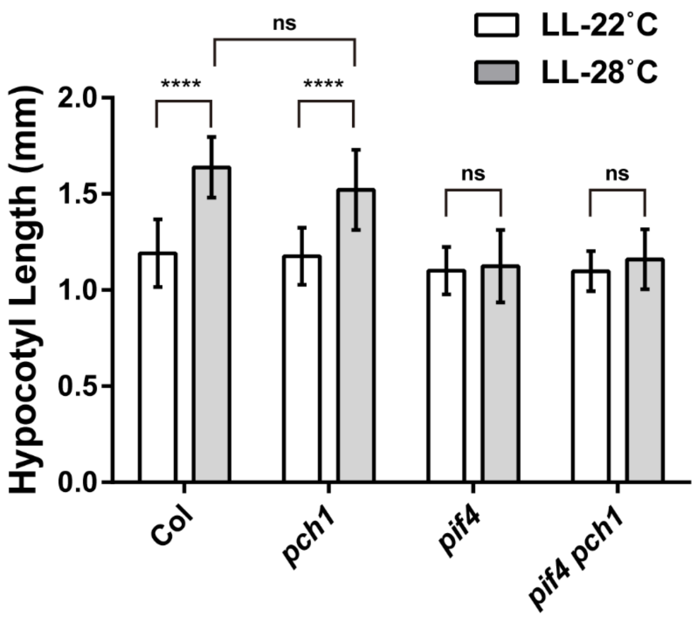


**Fig. S6.** Genetic interactions between *pif4* and *pch1* under constant light (LL) with different temperature (LL-22˚C and LL-28˚C) conditions in regulating hypocotyl elongation. Two-way ANOVA and Tukey's multiple comparisons tests were conducted (n = 20). Error bars = standard deviation. ns, not significant; ****, p<0.0001. The experiment in this figure was done twice with consistent results.

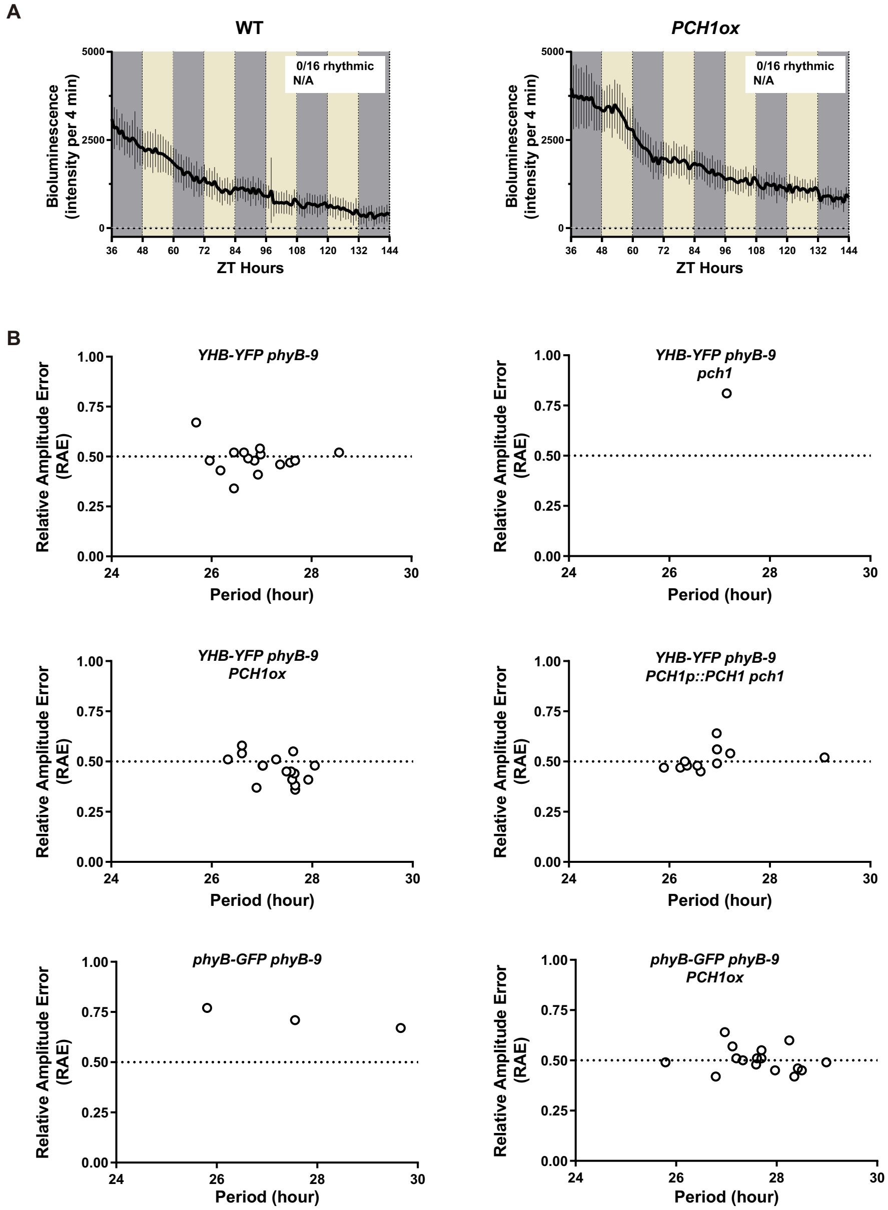


**Fig. S7.** Relative amplitude error **(**RAE) analysis of *YHB-YFP*/*phyB-GFP* plants with altered PCH1 levels.
(*A*) Bioluminescence intensity of the *CCA1::LUC* reporter in Col-0 and *PCH1ox* seedlings. Seedlings were treated and analyzed same as in **Fig. 4*B***.
(*B*) RAE analysis of *YHB-YFP phyB-9* and *phyB-GFP phyB-9* plants in **Fig. 4*B***. Dots for each genotype are seedlings exhibiting cycling bioluminescence of the *CCA1::LUC* reporter. RAE was plotted as a function of measured circadian period. The dash line at RAE = 0.5 serves as a cutoff and any point above this line is not considered reliably rhythmic (9).
All experiments in this figure were done twice with consistent results.

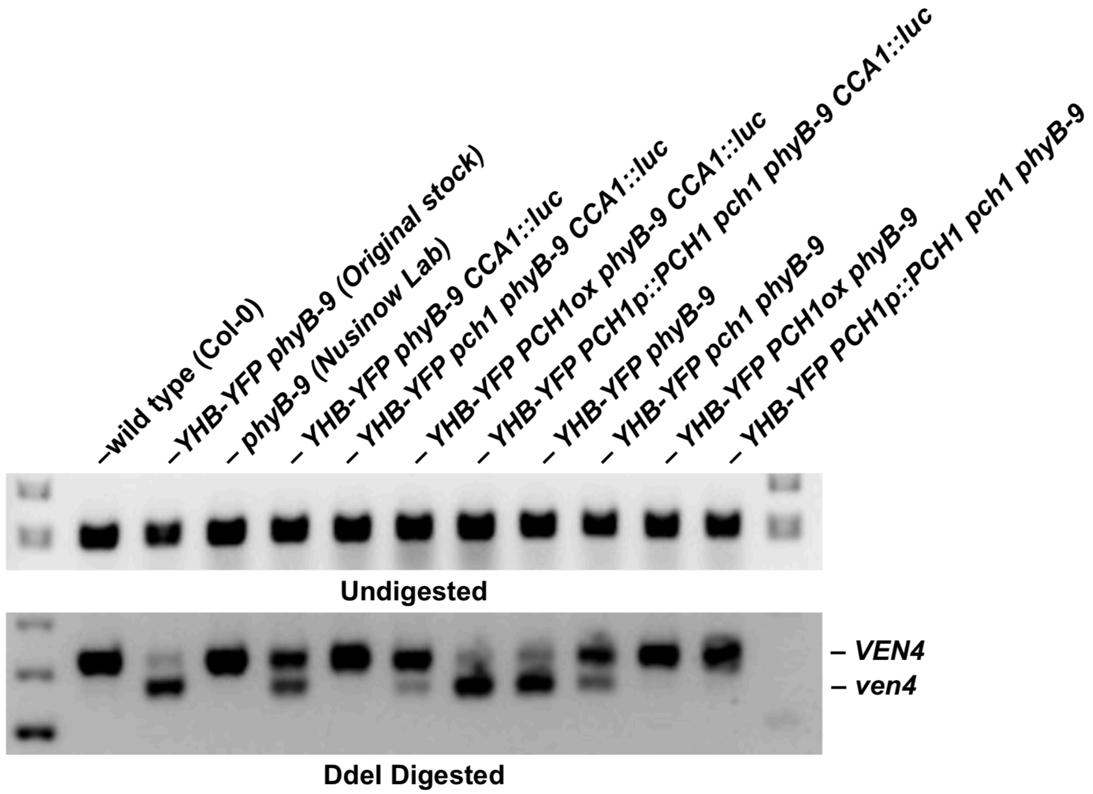


**Fig. S8.** Genotyping of VENOSA4 in the YHB lines used in this study.
Approximately 15 seedlings were collected from seed stocks and genotyped for *VEN4/ven4* as described in (10). Upper gel (1% agarose) uncut amplicons, lower gel (3% agarose) DdeI digested products. Genotyping indicated that *YHB-YFP phyB-9* plants used for crossing introduced *ven4* into the lines, however, segregation of *ven4* does not correlate with loss of function or rescue phenotypes observed.

**Table S1.** List of all primers used.

| **Primers used for cloning (without a stop codon)** [a] | | |
|---|---|---|
| Amplified Fragments | Forward primer (5'->3') | Reverse primer (5'->3') |
| *phyB-FL* | *CACCATGGTTTCCGGAGTCGGGGGTAG* | *ATATGGCATCATCAGCATCATGTC* |
| *phyB-PSM (AA 1~639)* | *CACCATGGTTTCCGGAGTCGGGGGTAG* | *ATCCCTACATGGCTGAACCACA* |
| *phyB-OPM (AA 640~1172)* [b] | *CACCATGGCGGGGGAACAGGGGATTGATGAG* | *CTTGTTTGCTGCAGCGAGTTC* |
| *phyB-PASII&HKR (AA 777~1172)* | *CACCATGGATAAGTTCATCAACATACAAGGAG* | *CTTGTTTGCTGCAGCGAGTTC* |
| *phyB-PASII (AA 777~930)* | *CACCATGGATAAGTTCATCAACATACAAGGAG* | *TTTTGCCTTCGTGAAACACTCTGTGTC* |

| **Primers used for cloning (with a stop codon)** [a] | | |
|---|---|---|
| Amplified Fragments | Forward primer (5'->3') | Reverse primer (5'->3') |
| *phyB-FL* | *CACCATGGTTTCCGGAGTCGGGGGTAG* | *CTAATATGGCATCATCAGCATCATGTC* |
| *phyB-PSM (AA 1~639)* | *CACCATGGTTTCCGGAGTCGGGGGTAG* | *CTAATCCCTACATGGCTGAACCACA* |
| *phyB-OPM (AA 640-1173)* [b] | *CACCATGGCGGGGGAACAGGGGATTGATGAG* | *CTACTTGTTTGCTGCAGCGAGTTC* |
| *phyB-PASII&HKR (AA 777~1172)* | *CACCATGGATAAGTTCATCAACATACAAGGAG* | *CTAATATGGCATCATCAGCATC* |
| *phyB-PASII (AA 777~930)* | *CACCATGGATAAGTTCATCAACATACAAGGAG* | *TTATTATTTTGCCTTCGTGAAACACTCTGTGTC* |

| **Primers for cloning phyB-OPM-ΔPASII** | | |
|---|---|---|
| Amplified Fragments | Forward primer (5'->3') | Reverse primer (5'->3') |
| *Step1:phyB-PASI* | CACCATGGCGGGGGAACAGGGGATTGATGAG | GCTCGGGATTCCTTGTATGTTGATGAACTTAT |
| *Step1: phyB-HKR-stop* | ATACAAGGAATCCCGAGCCCTGAGCTGCA | CTAATATGGCATCATCAGCATCATGTCA |
| *Step1: phyB-HKR-nonstop* | ATACAAGGAATCCCGAGCCCTGAGCTGCA | ATATGGCATCATCAGCATCATGTCAC |
| *Step2: phyB-OPM-ΔPASII-stop* | CACCATGGCGGGGGAACAGGGGATTGATGAG | CTAATATGGCATCATCAGCATCATGTCA |
| *Step2: phyB-OPM-ΔPASII-nonstop* | CACCATGGCGGGGGAACAGGGGATTGATGAG | ATATGGCATCATCAGCATCATGTCAC |

| **Primers for generating phyB$^{YH}$** | | |
|---|---|---|
| Amplified Fragments | Forward primer (5'->3') | Reverse primer (5'->3') |
| *phyB*$^{YH}$ | CGTGTTATGGTTCATAAGTTTCATG | CATGAAACTTATGAACCATAACACG |

| **Primers for genotyping** | | |
|---|---|---|
| Mutant name | for wild type PCR (5'->3') | for mutant PCR (5'->3') |
| *pch1* [b] | *TGTCAGGTATTTCGGTCCTTG (LP)* and *CACTTGCTTGATGCTCATGAG (RP)* | *AAGAACCGGCAAAGATACCAC (RP)* and *ATTTTGCCGATTTCGGAAC (LBb 1.3)* |
| *pif4* (*pif4-101*) [c] | *CTCGATTTCCGGTTATGG (SL42)* and *CAGACGGTTGATCATCTG (SL43)* | *GCATCTGAATTTCATAACCAATC (PD14)* and *CAGACGGTTGATCATCTG (SL43)* |
| *phyB-9* [b] | *GTGGAAGAAGCTCGACCAGGCTTTG* and *GTGTCTGCGTTCTCAAAACG*, cut with MnlI, *phyB-9* gives 167+18 bp bands, WT gives a 185 bp band. | |
| *Ven4* [d] | *TGTAATGTACGTGCTTAACTTTCTCT* and *TCAAGGAATGTGCAAACATA,* cut with DdeI, *ven4* gives 235 + 25 bp band, WT gives a 260 bp band. | |

| **Primers for making pCMX-GW-HA** | |
|---|---|
| Primer Name | Sequence (5'->3') |
| pDAN0820 [e] | TATCAGGATCGGTACCGCTAGCTACCCATACGATGTTCCAGATTACGCT |
| pDAN0821 [e] | TAGCTACCTAGCTAGTTAAGCGTAATCTGGAACATCGTATGGGTAGCTA |
| pDAN0830 | CAGATATCAGGATCGGTACCACAAGTTTGTACAAAAAAGCTGAACGA |
| pDAN0831 | GGTAGCTAGCGGTACTACCCTAGGTACAAGAAAGCTGAA |

[a] CACC (underscored) were added to forward primers for cloning into the pENTR/D-TOPO vectors

[b] reference (1)

[c] reference (11)

[d] reference (10)

[e] underscored nucleic acids encode the HA tag, with pDAN0820 being the forward primer and pDAN0821 being the reverse primer.

**References**


1. Huang H, *et al.* (2016) PCH1 integrates circadian and light-signaling pathways to control photoperiod-responsive growth in Arabidopsis. *eLife* 5:e13292.
2. Qiu Y, *et al.* (2015) HEMERA Couples the Proteolysis and Transcriptional Activity of PHYTOCHROME INTERACTING FACTORs in Arabidopsis Photomorphogenesis. *Plant Cell* 27(5):1409-1427.
3. Huang H, *et al.* (2016) Identification of Evening Complex Associated Proteins in Arabidopsis by Affinity Purification and Mass Spectrometry. *Mol Cell Proteomics* 15(1):201-217.
4. Burgie ES, *et al.* (2017) Photosensing and Thermosensing by Phytochrome B Require Both Proximal and Distal Allosteric Features within the Dimeric Photoreceptor. *Scientific Reports* 7(1):13648.
5. Feng CM, Qiu Y, Van Buskirk EK, Yang EJ, & Chen M (2014) Light-regulated gene repositioning in Arabidopsis. *Nat Commun* 5:3027.
6. Edelstein A, Amodaj N, Hoover K, Vale R, & Stuurman N (2010) Computer control of microscopes using microManager. *Curr Protoc Mol Biol* Chapter 14:Unit14 20.
7. Edelstein AD, *et al.* (2014) Advanced methods of microscope control using muManager software. *J Biol Methods* 1(2).
8. Plautz JD, *et al.* (1997) Quantitative Analysis of Drosophila period Gene Transcription in Living Animals. *Journal of Biological Rhythms* 12(3):204-217.
9. Harmer SL & Kay SA (2005) Positive and negative factors confer phase-specific circadian regulation of transcription in Arabidopsis. *Plant Cell* 17(7):1926-1940.
10. Yoshida Y, *et al.* (2018) The Arabidopsis phyB-9 Mutant Has a Second-Site Mutation in the VENOSA4 Gene That Alters Chloroplast Size, Photosynthetic Traits, and Leaf Growth. *Plant Physiol* 178(1):3-6.
11. Nusinow DA, *et al.* (2011) The ELF4-ELF3-LUX complex links the circadian clock to diurnal control of hypocotyl growth. *Nature* 475(7356):398-402.
